# TGFβ controls alveolar type 1 epithelial cell plasticity and alveolar matrisome gene transcription

**DOI:** 10.1101/2023.05.09.540035

**Authors:** Danielle A. Callaway, Ian J. Penkala, Su Zhou, Fabian Cardenas-Diaz, Apoorva Babu, Michael P. Morley, Mariana Lopes, Benjamin A. Garcia, Edward E. Morrisey

## Abstract

Premature birth disrupts normal lung development and places infants at risk for bronchopulmonary dysplasia (BPD), a disease increasing in incidence which disrupts lung health throughout the lifespan. The TGFβ superfamily has been implicated in BPD pathogenesis, however, what cell lineage it impacts remains unclear. We show that *Tgfbr2* is critical for AT1 cell fate maintenance and function. Loss of *Tgfbr2* in AT1 cells during late lung development leads to AT1-AT2 cell reprogramming and altered pulmonary architecture, which persists into adulthood. Restriction of fetal lung stretch and associated AT1 cell spreading through a model of oligohydramnios enhances AT1-AT2 reprogramming.

Transcriptomic and proteomic analysis reveal the necessity of *Tgfbr2* expression in AT1 cells for extracellular matrix production. Moreover, TGFβ signaling regulates integrin transcription to alter AT1 cell morphology, which further impacts ECM expression through changes in mechanotransduction. These data reveal the cell intrinsic necessity of TGFβ signaling in maintaining AT1 cell fate and reveal this cell lineage as a major orchestrator of the alveolar matrisome.

## INTRODUCTION

Bronchopulmonary dysplasia (BPD) is a lung disease that disproportionately affects infants born at less than 28 weeks gestation, the threshold for extreme prematurity^1^. Despite multiple medical advances including antenatal steroids, surfactant supplements, and improved mechanical ventilation strategies, disease incidence has increased as more infants are born at the limits of viability^2^. BPD is characterized as an arrest of alveolarization with enlarged airspaces and thickened septa indicative of a departure from the normal pulmonary developmental programming. This disease not only adversely impacts an infant’s immediate postnatal course during their stay in the neonatal intensive care unit, but it is a life-long disease with increased rehospitalization in the first two years of life^3, 4^, persistently impaired pulmonary function^5–7^, and increased risk of developing chronic obstructive pulmonary disease (COPD) as an adult^8^. Although the mechanisms for BPD pathogenesis are not fully known, one candidate that may play a role is transforming growth factor (TGFβ). Biologically active TGFβ in endotracheal aspirates is abundant in preterm infants and predicts the need for home oxygen therapy^9^. Furthermore, elevated levels of TGFβ ligand can be detected in bronchoalveolar lavage fluid from infants with BPD and correlates with disease severity^10^. However, the role of TGFβ signaling in the developing lung and how it differentially affects individual cellular niches, particularly the alveolar epithelium, is unclear.

The TGFβ signaling pathway is responsible for a multitude of critical cellular functions including apoptosis, proliferation, extracellular matrix (ECM) production, and mediating cell fate^11, 12^. Importantly, it is also a crucial relay for mechanical signaling within tissue as inactive TGFβ ligands are embedded within the ECM and become liberated and activated by cellular integrins through mechanical stress^13^. TGFβ receptors Tgfbr1 and Tgfbr2 as well as downstream Smad proteins exhibit dynamic temporal and spatial expression within the mouse lung with the highest levels of expression appearing during late lung development^12^. Rodent models of TGFβ deletion and overexpression have underscored its importance in regulating lung development^14^. Postnatal overexpression of TGFβ1 under an inducible Scgb1a1 promoter or through intranasal delivery of an adenoviral vector resulted in a BPD-like phenotype of alveolar enlargement^15, 16^. However, loss of Tgfbr2 or downstream components of the signaling pathway including Smad3 is not protective and instead similarly impairs alveologenesis^17–19^. Specifically, when Tgfbr2 is deleted from undifferentiated alveolar epithelial cells early during lung development (E6.5), the lungs demonstrated postnatal alveolar enlargement whereas deletion in the lung mesenchyme impaired branching morphogenesis^17^. Deletion of Tgfbr1 during lung endoderm development caused a blockade of secretory cell differentiation in mouse airways^20^. Our previous work has shown that TGFβ promotes *in vitro* cellular spreading of alveolar epithelial (AT1) cells ^21^. However, what role TGFβ plays in AT1 cell fate decisions or during pulmonary development is unclear.

In the current study, we show that *Tgfbr2* deletion in AT1 cells during late lung development increases AT1-AT2 cell reprogramming and induces a BPD-like phenotype of impaired alveolarization and increased septal thickness. A model of oligohydramnios, which predisposes to pulmonary hypoplasia and increases BPD risk in human neonates, impairs alveolar stretch and similarly drives AT1-AT2 cell reprogramming. Single-cell RNA sequencing reveals that expression of numerous genes associated with constituents of the pulmonary ECM and regulatory components, referred to as the pulmonary matrisome, are expressed in AT1 cells starting at the saccular stage through adulthood, reinforcing the AT1 cell as a node for matrisome production. RNA sequencing and proteomics of neonatal AT1 cells reveals that constituents of the pulmonary matrisome are downregulated upon loss of *Tgfbr2*. Moreover, TGFβ mediates AT1 cell integrin expression, which in turn affects cell size, morphology, and matrisome gene transcription. These studies reveal that TGFβ signaling is an important regulator of the pulmonary matrisome in AT1 cells, which controls sculpting of the developing alveolus to promote normal lung architecture and function.

## RESULTS

### Loss of Tgfbr2 results in increased AT1 cell reprogramming

To define the function of the main receptor of TGFβ signaling in AT1 cells, Tgfbr2 was deleted prenatally (Figure 1A-C) and postnatally (Figure 1D-H) using an AT1-specific tamoxifen-inducible transgenic mouse model (*Hopx^creERT^*^2^*:Tgfbr2:R26R^EYFP^*, hereafter referred to as *Tgfbr2^AT^*^1*-KO*^). Initially, pregnant dams were injected with tamoxifen at E15.5 followed by fetal lung harvest at E18.5, a point at which the majority of alveolar epithelial cells have undergone lineage specification (Fig. 1A)^25^. These experiments revealed that prenatal loss of *Tgfbr2* in AT1 cells yielded an increase in lineage-traced AT2 cells (Figure 1B-C). Similarly, postnatal tamoxifen injection at P0 followed by assessment at P5 and P42 also resulted in elevated AT1:AT2 cell reprogramming albeit to a greater extent at P42 suggesting that the influence of TGFβ on AT1 cell identity may be a permissive rather than direct effect (Figure 1D-F). Although TGFβ can regulate cellular proliferation, particularly in lung fibroblasts^39, 40^ and AT2 cells after injury ^41, 42^, proliferation of lineage-traced AT2 cells was unaffected by loss of *Tgfbr2* (Figure 1G-H). The overall numbers of AT1 and AT2 cells at P5 and P42 as a percent of total Nkx2-1-positive cells was also not significantly different (Figure S1A-C). There was no evidence of increased AT1 cell apoptosis after loss of *Tgfbr2* (Figure S1D).

**Figure 1.**
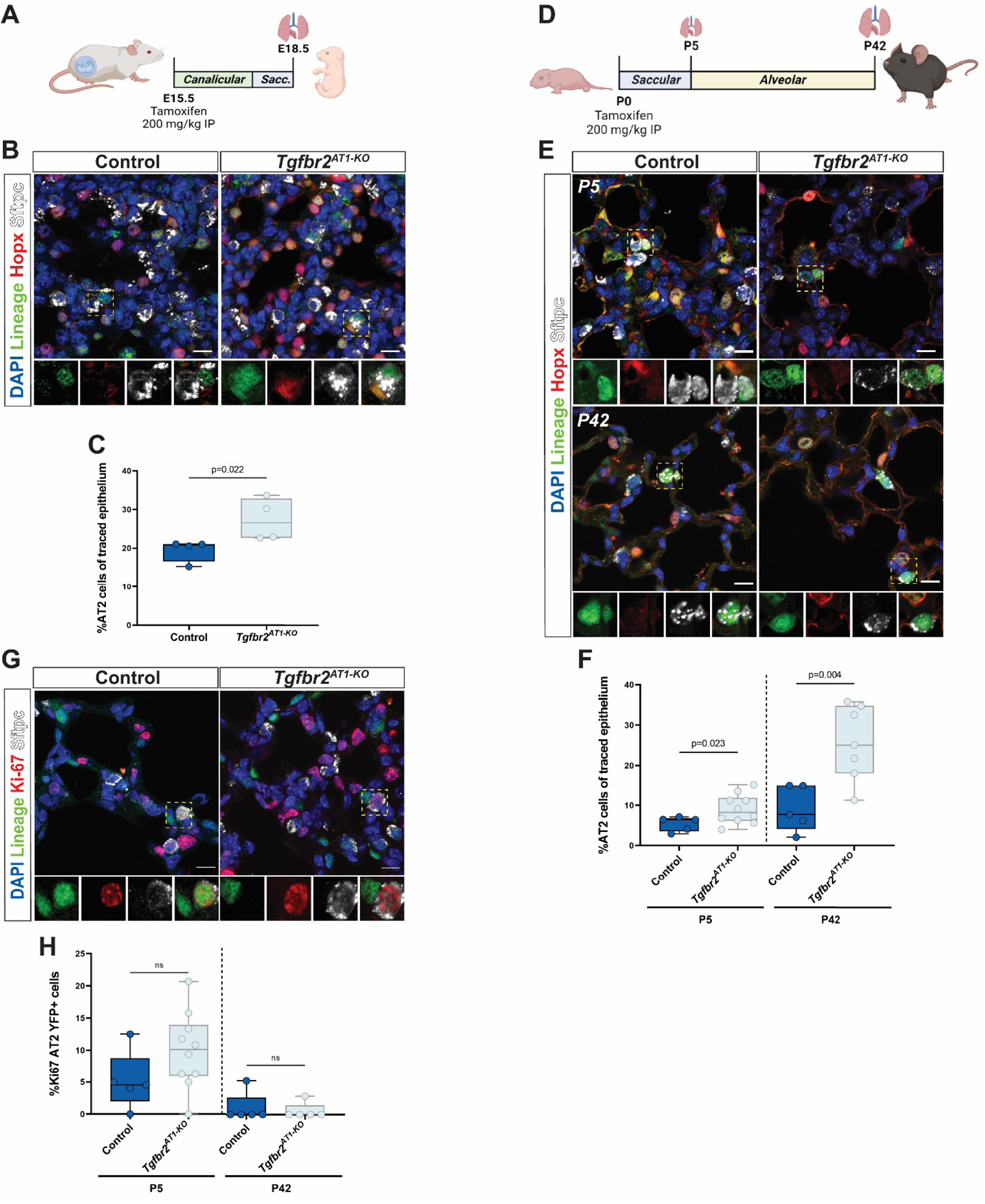
TGFβ is involved in regulating AT1 cell fate during prenatal and postnatal late lung development. A) Tamoxifen was delivered to control heterozygenous (*Tgfbr2^fl/+^*) littermates and knockout (*Tgfbr2^fl/fl^ or Tgfbr2^AT1-KO^*) through IP injection of the pregnant dam at E15.5 and lungs were harvested at E18.5. B) IHC for EYFP, HOPX, and SFTPC demonstrate increased AT1 reprogramming into AT2 cells after prenatal loss of *Tgfbr2*. The yellow dashed box denotes the magnified region shown below the image and separated by fluorescence channel. C) Quantification of lineage tracing in (B) denoting percent of cells that were EYFP+ and SFTPC+ by an unpaired two-tailed t-test (n=4 per group). D) In postnatal lineage-tracing experiments, control (*Hopx^creERT2^:R26R^EYFP^*) and *Tgfbr2^AT1-KO^* newborn pups (P0) were injected with tamoxifen and the lungs were harvested at P5 and P42. E) IHC for EYFP, HOPX, and SFTPC demonstrate increased AT1 reprogramming into AT2 cells after prenatal loss of *Tgfbr2* at both P5 (top) and P42 (bottom). The yellow dashed box denotes the magnified region shown below the image and separated by fluorescence channel. F) Quantification of lineage tracing in (E) denoting percent of cells that were EYFP+ and SFTPC+ at P5 (left) and P42 (right) by unpaired two-tailed t-tests with Welch’s correction (n=5-10 per group). G) IHC for EYFP, Ki67, and SFTPC indicate that there is no difference in lineage-traced AT2 cell proliferation across the groups. The yellow dashed box denotes the magnified region shown below the image and separated by fluorescence channel. H) Quantification of AT2 cell proliferation from (G) denoting percent of cells that were EYFP+, SFTPC+, and Ki67+ at P5 (left) and P42 (right) by unpaired two-tailed t-tests with Welch’s correction (n=5-10 per group). Each dot represents a single mouse. Scale bars denote 10 µm. ns = not significant. P values are denoted above the plots.

Previous work reported that loss of TGFβ signaling in mouse pulmonary epithelial cells alters pulmonary architecture to induce a BPD-like phenotype^17–19, 43^. Prenatal deletion of *Tgfbr2* in AT1 cells led to increased septal thickness (Figure 2A-B). Postnatal loss of *Tgfbr2* from birth through P5 and tracked through P42, a time-point at which alveologenesis has completed, resulted in alveolar simplification as demonstrated by increased mean linear intercept (MLI) as well as increased mean septal thickness at P5 (Figure 2C-E), indicating that AT1 specific loss of TGFβ during postnatal lung development has lasting effects on pulmonary architecture that persist into adulthood.

**Figure 2.**
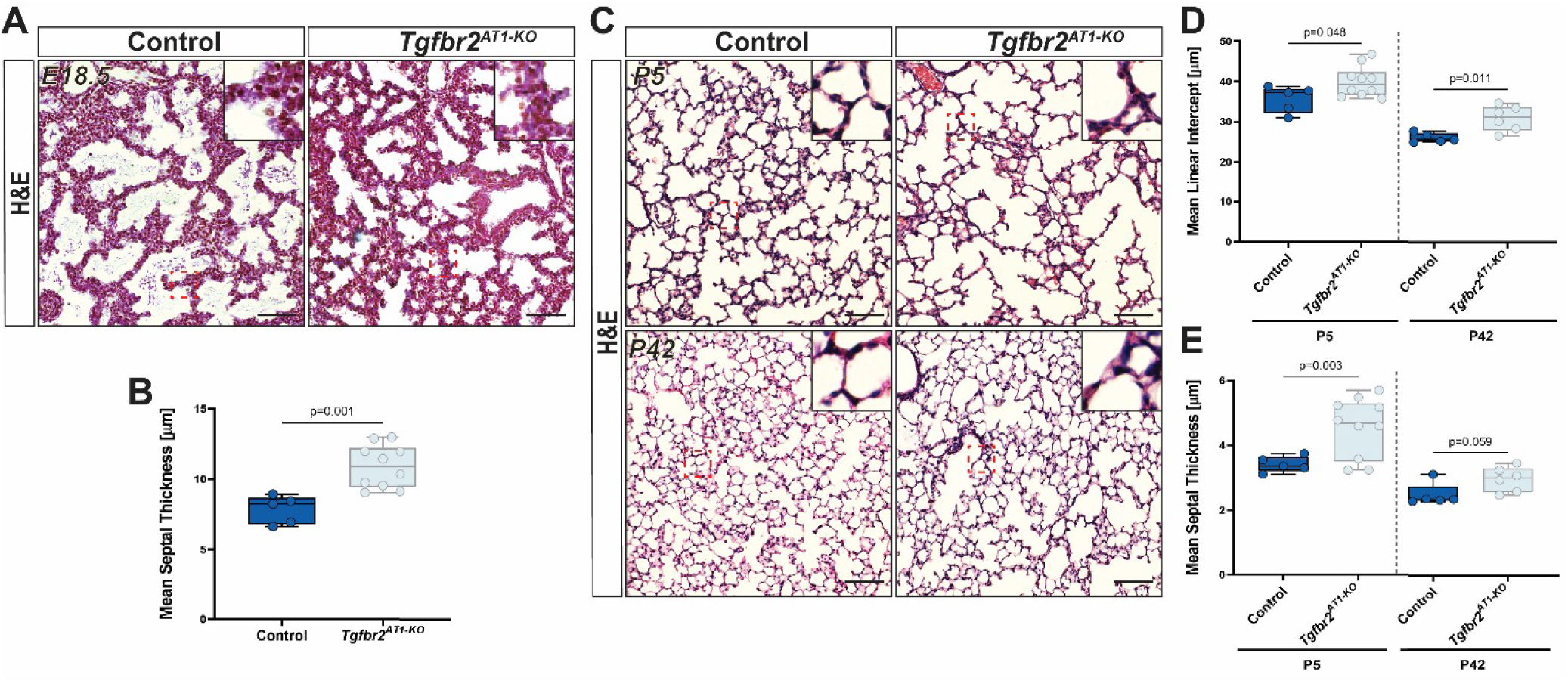
TGFβ is involved in maintaining normal pulmonary architecture during late lung development. A) H&E images of lungs from E18.5 mice following tamoxifen injection at E15.5 show that *Tgfbr2^AT1-KO^* embryonic lungs compared to heterozygous littermates demonstrate increased mean septal thickness. Red-dashed boxes indicate the zoomed area shown in the top-right of the image. B) Quantification of mean septal thickness (µm) from (A) by an unpaired two-tailed t-test (n=5-10 per group). C) H&E images of lungs from P5 (top) and P42 (bottom) mice following tamoxifen injection at P0 indicate that loss of TGFβ postnatally results in alveolar simplification and increased mean septal thickness. Red-dashed boxes indicate the zoomed area shown in the top-right of the image. D) Quantification of mean linear intercept (µm) and (E) mean septal thickness (µm) by unpaired two-tailed t-tests with Welch’s correction (n=5-10 per group). Each dot represents a single mouse. Scale bars denote 50 µm. P values are denoted above the plots.

### Lack of prenatal mechanical stretch promotes AT1 cellular reprogramming and AT2 cell fate specification

We have shown recently that maintenance of epithelial cell fate is intimately tied to mechanical signals whereby restriction of alveolar stretch in an adult lung deflation model led to reprogramming of AT1 cells into AT2 cells^44^. In the human neonate, prenatal reduction of amniotic fluid, termed oligohydramnios, can lead to pulmonary hypoplasia resulting in other respiratory-related morbidities including air-leak, pulmonary hypertension, and bronchopulmonary dysplasia^45^. Although the mechanism for how oligohydramnios directly leads to pulmonary hypoplasia is unclear, it is suspected to involve a restriction of thoracic expansion, thereby altering the intrapulmonary mechanical forces that are essential to distend the airways and promote normal lung maturation^46, 47^. A previous murine oligohydramnios model in which amniotic fluid was reduced at E15.5 followed by analysis at E18.5 resulted in reduced transcription of the AT1 cell-associated gene *Pdpn* and elevated expression of the AT2 cell marker *Sftpc* compared to non-treated littermates^29^. We performed a similar series of experiments with the inclusion of a lineage trace for AT1 cells (Figure 3A-B). Oligohydramnios resulted in increased mean septal thickness (Figure 3C-D) and increased total numbers of AT2 cells as a percentage of Nkx2.1+ epithelium (Figure 3E-F). Importantly, oligohydramnios resulted in increased AT1-AT2 cell reprogramming (Figure 3E-G), indicating that inhibiting essential mechanical forces during late pulmonary development is sufficient to alter epithelial cell fate in the fetal lung.

**Figure 3.**
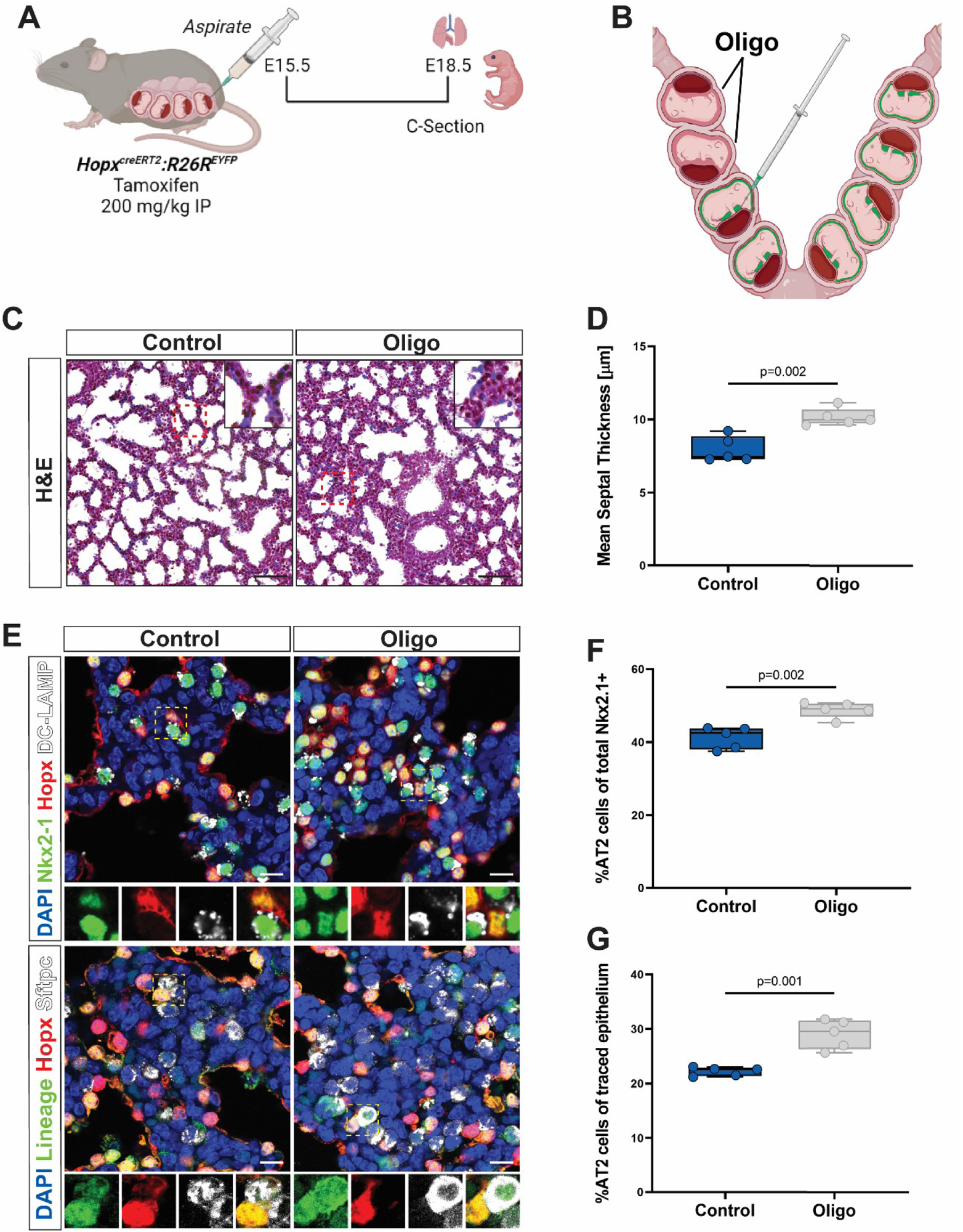
Oligohydramnios alters alveolar epithelial cell numbers through increased AT1:AT2 cell reprogramming. A) Oligohydramnios was induced through amniotic fluid reduction at E15.5 followed by injection of tamoxifen to enable lineage tracing. Lungs from the fetal mice were harvested at E18.5 for further analysis. B) Amniotic fluid (green) was aspirated from amniotic sacs in the right uterine horn. C) H&E images of control (left) and oligohydramnios (right) lungs reveal that lack of amniotic fluid leads to increased mean septal thickness. D) Quantification of mean linear intercept (µm) by an unpaired two-tailed t-test (n=5 per group). E) IHC for NKX2.1 (TTF-1), HOPX, and DC-LAMP (top) reveal oligohydramnios leads to increased numbers of AT2 cells. Lineage-tracing through IHC for EYFP, HOPX, and SFTPC demonstrate that greater AT2 cell numbers may be secondary to increased AT1:AT2 cell reprogramming. The yellow dashed box denotes the magnified region shown below the image and separated by fluorescence channel. F) Quantification of AT2 cell percentage of total NKX2.1+ cells and (G) lineage tracing by unpaired two-tailed t-tests (n=5 per group). Each dot represents a single mouse. Scale bars denote 50 µm for H&E and 10 µm for IHC. P values are denoted above the plots.

### AT1 cells are enriched in genes associated with the pulmonary matrisome and is controlled by TGFβ

To determine gene expression changes due to loss of *Tgfbr2* in AT1 cells, RNA-sequencing of sorted AT1 cells from neonatal *Tgfbr2^AT^*^1*-KO*^ mice and heterozygous littermates at P5 was performed. One of the most significantly downregulated genes associated with *Tgfbr2* loss was *P3h2*, a member of the prolyl 3-hydroxylase subfamily that controls post-translational 3-hydroxylation of proline residues on collagen IV, which is necessary for collagen cross-linking and stability^48^. Because TGFβ is a known modulator of both ECM production and maintenance in the lung^14, 49–51^, we interrogated the dataset for other matrisome-related genes^37^. We found that in addition to *P3h2*, an additional ECM regulator, *MMP14*, was also downregulated as well as the ECM glycoprotein *Igfbp7* and matrisome-associated secreted factors *Lgals1* and *Ctf1*. *Cxcl5*, a secreted factor involved in neutrophil recruitment after injury, was upregulated (Figure 4A-B). Evaluation of gene ontology pathway enrichment revealed that loss of *Tgfbr2* in AT1 cells during late lung development was associated with enrichment of cellular components of the ECM and genes involved in ECM binding (Figure 4C-D). We next assessed changes in the proteome by mass spectrometry analysis. These studies revealed downregulated expression of several ECM-related proteins including glycoproteins Lama3 and Igfbp7, proteoglycans Spock2 and Hspg2, ECM regulators Adam10 and Ctsh, and integrins Itga3 and Itgb1 (Figure 4E).

**Figure 4.**
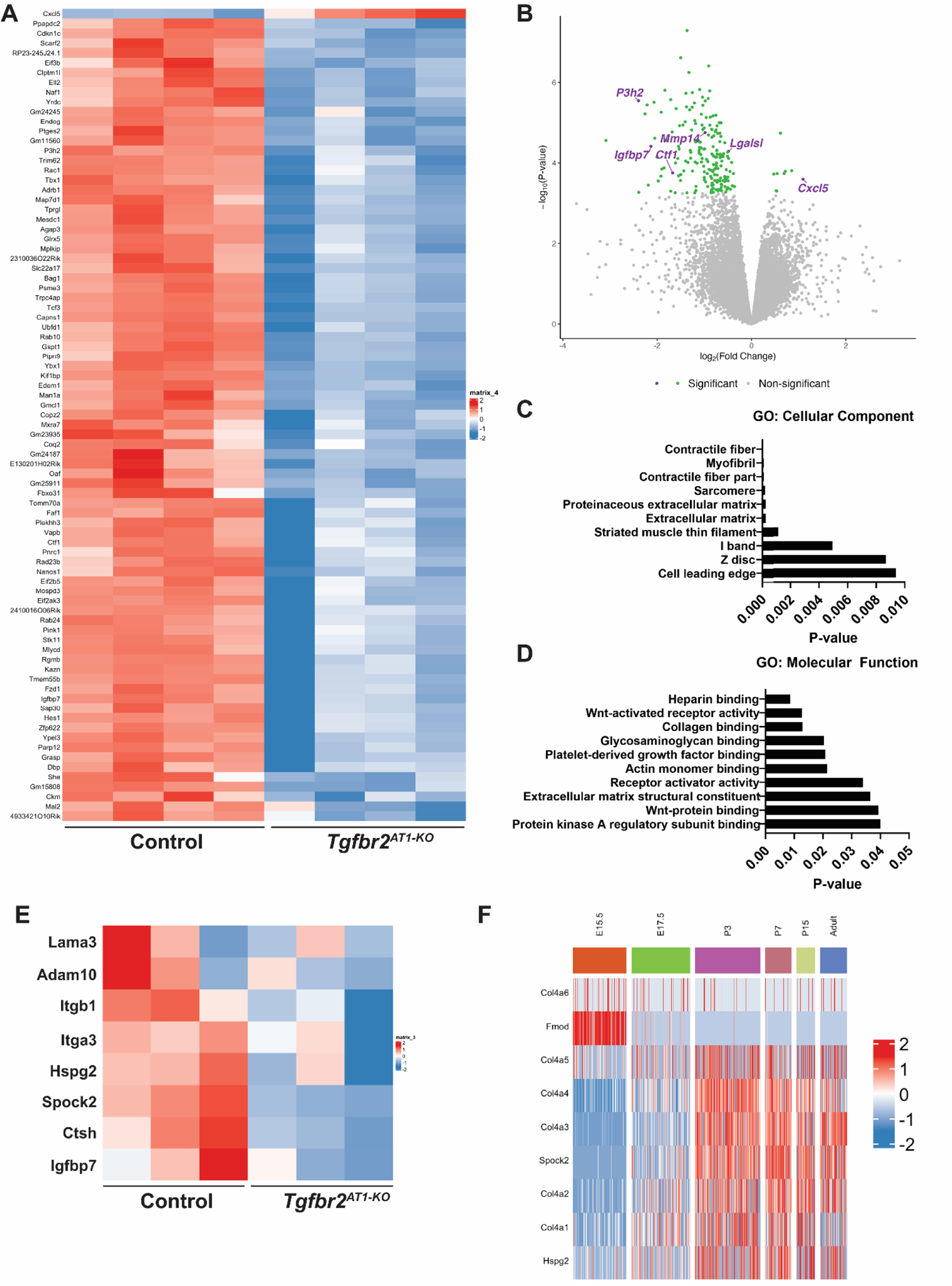
Transcriptomic and proteomic profiling reveal a role for TGFβ in regulating AT1 cell expression of the pulmonary matrisome. A) Heatmap of differentially expressed genes from AT1 cells of control heterozygous littermates (left) or *Tgfbr2^AT1-KO^*(right) mice at P5 (n=4). B) Volcano plot of differentially expressed genes from RNA-seq from (A) with ECM-related genes labeled reveal that several are significantly downregulated with loss of *Tgfbr2*. C) GO enrichment for cellular component and (D) molecular function of downregulated genes reveal several ECM-related terms. E) Heatmap of ECM-related proteins from proteomics results of control heterozygous littermates (left) or *Tgfbr2^AT1-KO^* (right) AT1 cells obtained at P5 (n=3). F) Evaluation of previously generated scRNA-seq data across development indicates that AT1-enriched collagen IV and proteoglycan genes become highly expressed in the postnatal period through adulthood indicating a role for AT1 cells in the active production and remodeling of the pulmonary matrisome across the lifespan.

Transcriptomic and epigenetic profiling of AT1 cells has revealed that they are highly enriched for pathways involving focal adhesion, integrin-mediated cell adhesion, and cytoskeleton regulation^26, 28^. Single-cell RNA sequencing of the developing lung has highlighted the importance of AT1 cells in transcription of ECM related genes including collagens, proteoglycans, and glycoproteins^28, 52^. *Col4a3* and *Col4a4* as well as Laminin-332 constituents *Lama3*, *Lamb3*, and *Lamc2* have higher expression in AT1 cells compared to other cell populations within the lung^52^. Using a previously generated single-cell sequencing (scRNA-seq) dataset, we compared other AT1-enriched core matrisome constituents across development including collagens and proteoglycans (Figure 4F) as well as glycoproteins (Figure S2A)^28, 37^. Most AT1 cell enriched matrisome-related genes become highly expressed at E17.5, corresponding with the start of the saccular stage of lung development. Nearly all of these genes remain highly expressed through adulthood indicating a life-long importance of AT1 cells as critical orchestrators for the production and maintenance of the pulmonary matrisome. Similarly, integrin expression in AT1 cells also increases starting at E17.5 and remains elevated through adulthood with the exception of the laminin-binding integrin Itga6, which appears to exhibit less expression postnatally (Figure S2B).

Because of the importance of core matrisome components including collagens, proteoglycans, and glycoproteins as well as ECM regulators to AT1 cell biology, we elected to expand our scope beyond what was seen in the informatics analysis to evaluate additional matrisome components in *Tgfbr2* deficient AT1 cells. The specific collagen and laminin subtypes were chosen due to their high expression in AT1 cells as depicted in scRNA-seq data^28, 52^. The collagen IV and laminin-332 genes were decreased in Tgfbr2^AT1-KO^ cells, although to a greater extent at P42 (Figures 5A-F, S3A). Corresponding *Col4a3* and *Col4a4* RNA-scope indicated the relative specificity for each transcript in the *Hopx*^+^ labeled cells compared to *Lamb3*, which is present in several other non-labeled cells in agreement with data from scRNA-seq (Figures 5A, C, E). The remaining genes were chosen due to their presence in the RNA-sequencing and/or proteomics data and included the glycoprotein *Igfbp7*, proteoglycan *Hspg2*, and ECM regulator *P3H2*. *Plod2*, a gene that is involved in collagen hydroxylation and crosslinking, was chosen as it is known to be regulated by TGFβ^53^. Importantly, expression of each of these genes was decreased in the Tgfbr2-deficient AT1 cells from neonates and adults (Figures 5G-H, S3B-D).

**Figure 5.**
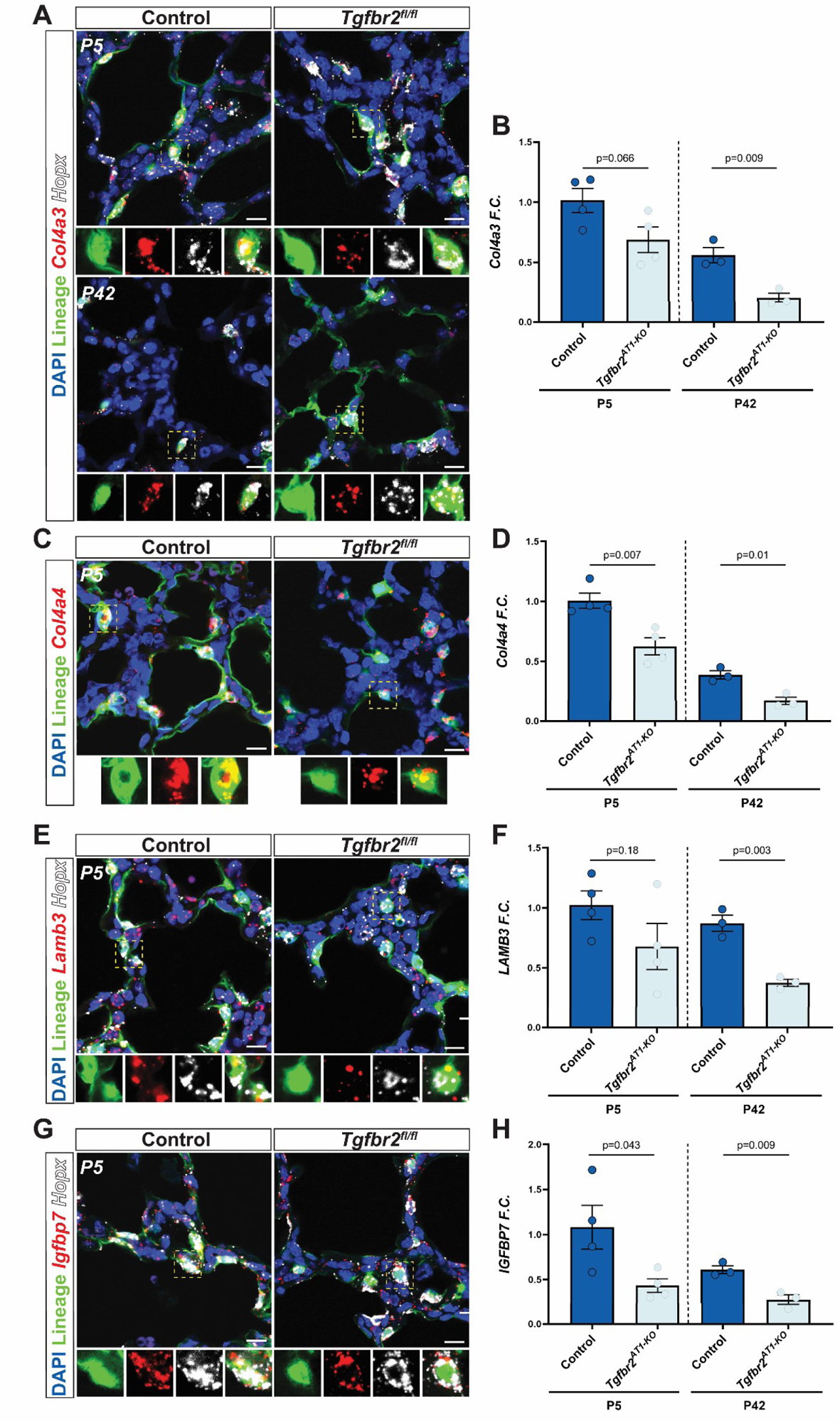
Loss of TGFβ at birth perturbs AT1-mediated matrisome expression through adulthood. A) RNA FISH for *Col4a3* and *Hopx* with IHC for lineage-labeled cells at P5 (top) and P42 (bottom). The yellow dashed box denotes the magnified region shown below the image and separated by fluorescence channel. B) *Tgfbr2^AT1-KO^* AT1 cells exhibit decreased RNA transcript expression for several AT1 cell-enriched core matrisome constituents including collagens *Col4a3* and (D) *Col4a4* and glycoproteins (F) *Lamb3* and (H) *Igfbp7 at* P5 that persists to P42 (n=3-4, unpaired two-tailed t-test with Welch’s correction). D) RNA FISH of P5 lungs for *Col4a4* with IHC for lineage-labeled cells. The yellow dashed box denotes the magnified region shown below the image and separated by fluorescence channel. E) RNA FISH for *Hopx* and glycoproteins *Lamb3* and (G) *Igfbp7* with IHC for lineage-labeled cells. The yellow dashed box denotes the magnified region shown below the image and separated by fluorescence channel. Each dot represents a single mouse. Scale bars denote 10 µm. P values are denoted above the plots.

### TGFβ affects AT1 cell spreading and morphology through transcriptional regulation of integrins

Prior work from our lab has established a role for TGFβ in promoting AT1 cell spreading^54^. We wanted to evaluate the consequence of increased or inhibited TGFβ signaling on AT1 cell spreading as well as their effects on matrisome transcription. We isolated AT1 cells from neonatal mice and generated primary AT1 cell cultures, which were treated with TGFβ1 ligand or the TGFβ inhibitor SB431542 (Figure 6A). Cell size was quantified every other day of culture up to 6 days and RNA was collected for analysis. The cultured cells expressed the AT1 cell marker Ager and gradually spread through the course of the experiment (Figures 6B-C). Treatment with SB431542 decreased the extent of AT1 cell spreading while exogenous TGFβ1 ligand had no significant affect (Figure 6C). In contrast, there was a difference in cell morphology upon treatment with exogenous TGFβ1 (Figures 6B, D). While the cells treated with SB431542 spread less but acquired a more stellate appearance, the cells treated with exogenous TGFβ1 appeared rounder and demonstrated more uniform spreading compared to the controls (Figures 6B, D). This alteration in cell shape was visual findings were evaluated by measuring cell roundness with the Fiji imaging software package, which confirmed that TGFβ1 treated cells had a higher roundness score compared to the control and inhibitor-treated cells (Figure 6B and D).

**Figure 6.**
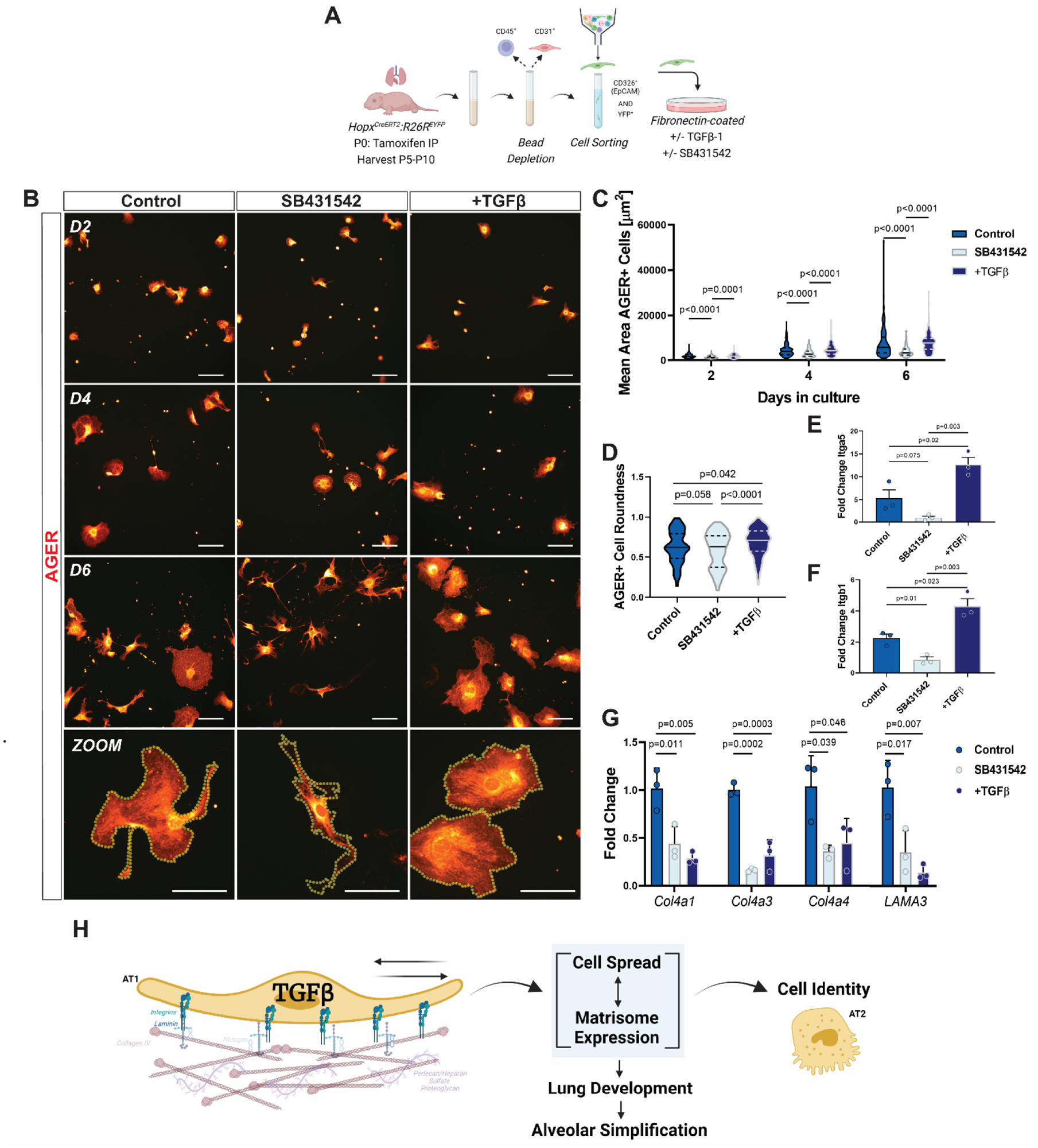
TGFβ-mediated integrin binding regulates AT1 cell size, morphology, and ECM expression. A) To obtain AT1 cells for culture, P5-P10 pup lungs were obtained after lineage labeling with tamoxifen at P0. Whole lung cell suspensions were obtained using a dispase, DNase, and collagenase digestion buffer after which the epithelial cell population was enriched by depleting the CD45+ and CD31+ population with magnetic beads. The remaining cells were fluorescently sorted with FACS to obtain a CD326+ and YFP+ suspension. The cells were plated onto fibronectin-coated plates with or without TGFβ-1 ligand (7.5 ng/mL) or the TGFβ inhibitor SB431542 (10µM) and evaluated at days 2, 4, and 6. B) ICC of RAGE+ (AGER+) cells treated with TGFβ ligand or inhibitor in culture at days 2, 4, and 6 with a zoomed image of cells at day 6 appearing at the bottom. Exogenous TGFβ resulted in a more equal spread of the cell whereas treatment with inhibitor led to decreased cell size. Scale bars denote 100 µm. C) Quantification of mean AGER+ cell area depicted in (B) by one-way ANOVA with Tukey’s multiple comparisons (n=135-231). D) Quantification of mean cell roundness at day 6 depicted in (B) by one-way ANOVA with Tukey’s multiple comparisons (n=148-167). E) Quantification of transcript expression levels (fold change compared to GAPDH) of the fibronectin-binding integrins *Itga5* and (F) *Itgb1* indicate that TGFβ mediates their expression (n=3 per group, one-way ANOVA with Tukey’s multiple comparisons). H) Quantification of transcript expression levels (fold change compared to GAPDH) for several basement membrane components including collagen IV subtypes *Col4a1*, *Col4a3*, and *Col4a4* and laminin-332 constituents *Lama3* and *Lamb3* indicate that altering homeostatic TGFβ signaling reduces expression of several AT1-specific ECM genes (n=3 per group, one-way ANOVA with Tukey’s multiple comparisons). I) Representative schematic of findings indicating that TGFβ regulates integrin expression to guide ECM binding and cellular spread, which affects cell identity and matrisome expression and impacts lung development.

We examined whether expression of integrins and AT1 matrisome components identified in our *in vivo* loss of function studies were altered in cultured AT1 cells treated with TGFβ1 ligand or inhibitor. Treatment with TGFβ1 significantly increased both *Itga5* and *Itgb1* expression compared to the control and inhibitor-treated groups while inhibition decreased their expression (Figures 6E-F). Loss of TGFβ signaling in cultured AT1 cells led to decreased expression of *Col4a1*, *Col4a3*, *Col4a4*, *Lama3*, and *Lamb3* (Figure 6G). Surprisingly, addition of exogenous TGFβ1 led to a similar decrease in expression of these matrisome components (Figure 6G). These data suggest that AT1 cells respond to TGFβ activation via a negative feedback loop, which suppresses expression of some matrisome components, likely to dampen excessive ECM deposition. Together, these data support the important function of TGFβ as a mediator of AT1 cellular spreading through integrin-binding to the ECM. AT1 spreading and its associated changes in mechanotransduction are intimately tied to AT1 cell identity and the ability of these cells to function as a node for matrisome expression in the lung (Figure 6H).

## DISCUSSION

In this study, we demonstrated the importance of TGFβ signaling in AT1 cell fate, which when disrupted, leads to a BPD-like phenotype with enlarged alveoli and increased septal thickness. Using a model of mouse oligohydramnios, which limits lung prenatal cyclical stretch, we show that the fate of prenatal AT1 cells is exquisitely sensitive to the natural breathing movements exhibited by the mammalian lung prior to birth. TGFβ signaling mediates the expression of multiple members of the pulmonary matrisome in AT1 cells including integrins, which are necessary to guide and sculpt the emerging alveolus during late lung development. Ex vivo modeling of AT1 cell spreading shows that TGFβ signaling impacts both AT1 cell morphology and overall extent of spreading, through integrin-mediated binding to the ECM, which is sensitive to negative feedback from excessive TGFβ activity. These data highlight AT1 cells as a nodal organizing center for sculpting and remodeling the lung alveolus during late development through TGFβ-mediated expression of the pulmonary matrisome.

RNA sequencing shows that AT1 cells are enriched for pathways involving focal adhesion, the actin cytoskeleton, ECM-receptor interactions, Hippo signaling, and TGFβ signaling, which are all related to mechanotransduction ^26, 44^. Neonatal loss of YAP/TAZ in AT1 cells results in a striking increase in AT1-AT2 cellular reprogramming and alveolar simplification^26^. Further, *in vivo* loss of interactions between the cytoskeleton and ECM through Cdc42 and Ptk2 in adult AT1 cells promotes AT1-AT2 cell reprogramming^44^. Conversely, physically constraining AT2 cells from spreading *in vitro* inhibits their AT2-AT1 differentiation^44^. Blockade of stretch during breathing through bronchial ligation also results in AT1-AT2 cell reprogramming, underscoring the essential role of mechanical signaling to maintain AT1 cell fate^44^. Impaired integrin expression in the present *Tgfbr2* loss of function study alters AT1 cell spreading ability, which affects expression of basement membrane constituents as well as enzymes for post-translational processing to ensure normal morphology. These changes further impair normal AT1 cell spreading, which promotes cellular reprogramming and a greater exacerbation of altered matrisome maintenance, which can lead to pathologic alveolarization and defects in secondary septation. These data support the emerging concept that AT1 cells are an organizing node for promoting and maintaining lung alveolar architecture.

In other organ systems including bone, which is under constant mechanical load, mechanotransduction is intimately tied to ECM expression in order to provide the necessary structure and support^55^. The lung, which is also exposed to continuous mechanical stress through cyclical breathing, likely regulates ECM production through constant sensing of mechanical strain in the various tissue niches. While several studies have explored ECM remodeling within the lung and the role of TGFβ, these have focused predominantly on the pro-fibrotic response of mesenchymal cells and AT2 cells in disease states such as cancer and idiopathic pulmonary fibrosis (IPF)^19, 47, 56, 57^. The novel role for AT1 cells as critical hubs for expression of core matrisome constituents, regulatory enzymes, and secreted factors during development has recently been described^28, 52^. AT1 cells produce constituents of the basement membrane including collagen IV, laminins, and other glycoproteins and proteoglycans that are important to maintain the structure and function of the alveolar-capillary barrier. Some components including *Col4a3, Col4a4,* and *Col4a5*, which make up the collagen IV α3α4α5 triple helix, and laminin-332 exhibit the greatest expression in AT1 cells compared to other cell types within the lung^52^. The collagen IV α3α4α5 isoform is thought to be stiffer owing to an increased number of crosslinks^58^. Furthermore, the collagen IV α3α4α5 isoform in the adult renal glomerular basement membrane (GBM) has been explored due to its role in Alport disease in which mutations in this isoform lead to progressive renal failure^59^. Goodpasture’s syndrome, another progressive renal disease in which antibodies against Col4a3 attack the basement membrane, results in pulmonary manifestations including hemoptysis and pulmonary hemorrhage^60^. Therefore, production and maintenance of the alveolar basement membrane by AT1 cells likely serves a critically important role to maintain the integrity of the alveolar-capillary barrier and promote cell spreading during alveologenesis to increase surface area for gas exchange. Beyond matrisome constituents, AT1 cells also influence neighboring cells through paracrine signaling of matrisome-related secreted factors that likely further guide pulmonary development and remodeling. Given the focal enrichment of AT1 matrisome expression in the adult, it is likely that the AT1 cell plays a key role in maintaining ECM-driven alveolar architecture maintenance and also may play an important role in fibrotic diseases of the lung such as IPF.

The lung is an organ under constant physical strain during normal breathing motions and displays remarkable regenerative capacity when challenged by injury. Late lung development marks a time of vulnerability in which the lung may be required to deviate from normal developmental programming and simultaneously respond to additional insults. Although the majority of injury models in neonates and adults involve infection, fibrosis, or hyperoxia, less is known regarding the effects of altered biophysical force. However, the potential clinical implications are important. There are many congenital anomalies that induce pulmonary hypoplasia including congenital diaphragmatic hernia, giant omphalocele, and renal dysplasias. Additionally, loss of amniotic fluid from preterm premature rupture of membranes and subsequent premature birth is a risk factor for BPD. Conversely, elevated biophysical force through exposure to mechanical ventilation induces TGFβ signaling, leading to disordered collagen, which can influence BPD pathogenesis in preterm infants. Future studies on the impact of mechanical stress on AT1-mediated matrisome maintenance will likely identify additional pathways that drive aberrant remodeling and intercellular communication during development and predisposes patients to diseases such as BPD and pulmonary hypoplasia.

## MATERIALS AND METHODS

### Animals

*Hopx^CreERT^*^2^ [Jackson Labs #017606]^22^, *R26R^EYFP^* (Jackson Labs #007903)^23^, and *Tgbr2^fl/fl^* (Jackson Labs #012603)^24^ have been previously described. All mice were maintained on a mixed background (C57BL/6 and CD-1) and all experiments were performed with at least n=3 per group of mixed sex. Controls were either heterozygous littermates or mice without the floxed allele as indicated in the text and figure legends.

### Lineage Tracing

Neonatal lineage tracing experiments were performed as previously described^25–28^. In studies of late lung development, pregnant dams at E15.5 gestation or newborn pups at P0 were injected intraperitoneally with a single 200 mg/kg dose of a solution containing tamoxifen (Sigma-Aldrich, T56480, 10% ethanol, and 90% corn oil (Sigma-Aldrich, C8267).

### Oligohydramnios Model

Amniotic fluid removal of pregnant dams at E15.5 to induce oligohydramnios and pulmonary hypoplasia of the embryos was performed as described in previous publications^29, 30^. Pregnant mice were anesthetized with isoflurane and prepared in sterile fashion. A longitudinal incision was made along the abdomen revealing the embryos and amniotic fluid was removed from embryos of the right uterine horn while those of the left remained untouched to serve as littermate controls. The mother was sutured closed and 200 mg/kg of tamoxifen was delivered intraperitoneally for lineage tracing experiments. At E18.5 the mother was euthanized with CO_2_ and the lungs of the progeny were removed for analysis.

### AT1 cell isolation from lungs

Lungs from tamoxifen-treated animals were removed and digested in a solution containing collagenase, DNase, and dispase as previously described^26, 28^. Red blood cells were lysed using ACK buffer and cells were then stained with EpCAM-PE-Cy7 (eBioscience, G8.8, 1:200), CD31-APC (eBioscience, 390, 1:200), and CD45-APC (eBioscience, 30-F11, 1:200). The CD45+ and CD31+ stained-cells were depleted on MACS LS columns after incubation with anti-APC magnetic beads (Miltenyi biotech) leaving an enriched epithelial cell population. The cell-containing solution was then sorted with a BD FACS Jazz cell sorter (Becton Dickinson) for YFP+ and CD326+ (EpCAM) double-positive cells to obtain lineage-traced AT1 cells for further analysis including RNA sequencing, proteomics, qRT-PCR, and *in vitro* studies.

### Histology and Alveolar Measurements

Animals were euthanized with CO_2_ and a thoracotomy was done to allow access to the lungs. The heart was perfused with cold PBS followed by inflation of the lungs with 2% paraformaldehyde at 25 cm H_2_O. The lungs were fixed overnight in 2% paraformaldehyde after which they were embedded in paraffin and sectioned at 6µm thickness. For alveolar morphology analysis, slides were stained with hematoxylin and eosin (H&E) per standard protocol. Images were obtained with a Nikon Eclipse 80i microscope or EVOS FL Auto2 Imaging System. Mean linear intercept (MLI) was performed with Fiji software across 10 images at 20x magnification per sample, counting the number of intercepts across 400 µm on 6 horizontal lines. Mean septal thickness was performed by measuring septal thickness in Fiji across 20 septa per image and 10 images per sample to yield 200 individual measurements that were averaged.

### Immunohistochemistry and RNA fluorescence in situ hybridization (FISH)

Immunohistochemistry was performed with the following antibodies: Cleaved Caspase 3 (rabbit, R&D Systems, MAB835, 1:100), Hopx (mouse, Santa Cruz, sc-398703 1:00), GFP (chicken, Aves Labs, GFP-1020, 1:200 or goat, Abcam, ab6673, 1:200), Ki67 (mouse, BD Biosciences, 550609, 1:200), Lamp3 (DC-Lamp) (rat, Novus, DDX0191P-100, 1:100), Nkx2.1 (Ttf1) (rabbit, Abcam, ab76013, 1:50), and Sftpc, (rabbit, Millipore Sigma, AB3786, 1:100). Slides were mounted using Slowfade Diamond Antifade Mountant (Invitrogen, catalog # S36972). RNA FISH was performed with RNAscope according the manufacturer’s instructions with the following probes: Mm-Col4a4-C2 (#1078041-C2), mm-Col4a3 (#544361), mm-Hopx-C2 (#405161-C2), mm-Igfbp7 (#425741), Mm-Lamb3 (#552161), and mm-Sftpc-C2 (#314101-C2). Fluorescent imaging was performed on a Zeiss LSM 710 confocal microscope with a 40x water immersion objective and processed using Fiji software^31^. For cellular quantification, at least 5 z-stacked images or 200 cells in total were assessed per sample.

### RNA sequencing (RNA-seq)

AT1 cells were isolated from neonatal mouse lungs at P5 as described. RNA isolation was done with the PureLink RNA Micro kit (ThermoFisher) and library preparation was performed with the NEBNext Single Cell/Low Input RNA Library Prep Kit for Illumina (New England Biolabs, catalog # E6420) following the manufacturer’s instructions. Libraries were sequenced using the Illumina HiSeq. Analysis was performed as previously described^32, 33^. In brief, the FastQC program was used to assess quality of Fastq files before aligning to the mouse reference genome (GRCm39) using the STAR aligner^34^. The MarkDuplicates program from Picard tools was used to flag duplicate reads and duplicates were excluded before computing per gene read counts for Ensembl (v104) gene annotations, using the Rsubread R package. Gene counts were normalized with the TMM method in the edge R package, and genes with 25% of samples with a counts per million (CPM) < 1 were deemed low expressed and removed^35^. Expression levels were transformed using VOOM in the limma R package. The limma R package was used to generate a linear model to perform differential gene expression analysis using linear models^36^. Given the small sample size of the experiment, we used the empirical Bayes procedure as implemented in limma to adjust the linear model fit and to calculate p values. These p values were adjusted for multiple comparisons using the Benjamini-Hochberg procedure. Plots were generated in R and GO enrichment was performed using the GAGE R package. The time course of single-cell RNA sequencing (sc-RNAseq) genes in AT1 cells was derived from previously generated data^28^ and heatmaps were generated in R. Matrisome-specific genes were derived from the Matrisome Project (https://sites.google.com/uic.edu/matrisome/home)^37^.

### Proteomics

Cells were lysed for 5 minutes on ice in 20 μL of lysis buffer containing 8 M urea, 0.1 M sodium chloride, and 50 mM Tris, pH 8.0, supplemented with phosphatase and protease inhibitors. Samples were centrifuged at 17900g for 5 minutes and the supernatant containing proteins was transferred to a new tube. Proteins were reduced using 5 mM DTT for 1 h at room temperature, and then alkylated with 20 mM iodoacetamide (IAA) in the dark for 30 min at room temperature. After that, samples were diluted with 4 volumes of 0.1 M ammonium bicarbonate and digested with 0.5 µg of trypsin overnight at room temperature. Samples were desalted and resuspended in 10 µL 0.1% formic acid for analysis (5 μL injections) by nLC-MS/MS, composed of an EASY-nLC 1000 (Thermo Scientific) coupled to a Q Exactive Orbitrap mass spectrometer (Thermo Scientific) operating in data-dependent acquisition (DDA) mode. Chromatography was performed at a flow rate of 300 nL/min over nano-columns (75 μm ID x 25 cm) packed with Reprosil-Pur C18-AQ (3 μm, Dr. Maisch GmbH). Water and 80% acetonitrile, both containing 0.1% formic acid, served as solvents A and B, respectively. The gradient consisted of 2–30% solvent B for 72 min, 30–60% B for 34 min, 60–90% B for 2 min, and 10 min of isocratic flow at 90% B. A full MS scan was acquired over 300–1400 m/z in the Orbitrap in in profile mode with a resolution of 70K, AGC target of 1e6, and maximum injection time of 100 ms. The top 15 most intense ions (charge state 2+ through 6+) were selected for MS/MS by high-energy collision dissociation (HCD) at 30 NCE, and fragmentation spectra were acquired in the Orbitrap in centroid mode with a resolution of 17.5K, AGC target of 1e5, isolation window of 2 m/z and maximum injection time of 50 ms. Dynamic exclusion of 40 s was used. Data was processed in Proteome Discoverer 2.2 (Thermo Scientific) using the Sequest-HT node to search MS/MS spectra against the UniProtKB/SwissProt (organism: Mus musculus) database along with a contaminant database. Trypsin was selected as the protease with a maximum of 2 missed cleavages. For the proteome data, the search parameters were as follows: 10 ppm precursor ion mass tolerance; 0.05 Da fragment ion mass tolerance; carbamidomethylation (+57.021 Da to Cys) as a static modification; and oxidation (+15.995 Da to Met) and acetylation (+42.011 Da to protein N-terminus) as variable modifications. The Percolator node was used with default parameters and data were filtered for < 1% FDR at the peptide level.

### Quantitative RT-PCR

Following fluorescent sorting of AT1 cells from mouse lungs, RNA isolation was performed with the PureLink RNA Micro kit (ThermoFisher). The SuperScript IV First-Strand synthesis system (Invitrogen) was used to generate cDNA and real-time qPCR was performed with Power SYBR Green Master Mix (ThermoFisher) on a QuantStudio 7 PCR System (Applied Biosystems).

### AT1 cell culture model, immunohistochemistry, and morphology analysis

Lineage-traced AT1 cells were obtained following FACS and were then placed into an organoid growth medium containing DMEM F12 and growth factors including bovine pituitary extract, cholera toxin, FBS, gentamicin, retinoic acid, insulin, transferrin, and human epithelial growth factor as previously described^27, 38^. A total of 25,000 cells were plated onto 24-well plates that were coated with fibronectin (5 µg/cm^2^, ThermoFisher). To evaluate the effects of exogenous TGFβ ligand or inhibition of TGFβ signaling, TGFβ1 (7.5 ng/mL, Peprotech) or SB 431542 (10 µM, Abcam) were added at the time of plating. Cells were fixed in 2% PFA at days 2, 4, and 6 or collected in RNA lysis buffer. After fixation, cells were stained with anti-RAGE antibody (rat, R&D, MAB1179, 1:100) and the entire well was imaged with the EVOS FL Auto2 Imaging System at 20x magnification. Only cells that were entirely within the image frame and were not adhered to an adjacent cell were assessed. Using Fiji software, cell area and roundness were quantified.

### Statistics

All statistics were performed with GraphPad Prism8 or in R Studio. Number of replicates and statistical test used is described in the figure legend. P values or “ns” (not significant) are denoted within the graph with significance determined by p<0.05.

### Study Approval

All studies were approved and performed in accordance with the University of Pennsylvania Institutional Use and Animal Care Committee.

### Data Availability

Sequencing data generated in this study has been deposited to GEO with accession number GSE230268. Proteomics data is available at MassIVE with accession number MSV000091803.

## AUTHOR CONTRIBUTIONS

D.A.C contributed to designing research studies, conducting experiments, acquiring data, analyzing data, and writing the manuscript. I.J.P designed research studies, conducted experiments, acquired data, and analyzed data. S.Z designed and conducted experiments. F.C designed research studies and analyzed data. A.B and M.P.M were involved in population RNA sequencing study design, data acquisition, data analysis, and figure generation. M.L and B.A.G were involved in proteomics study design, data acquisition, and data analysis. E.E.M contributed to designing research studies and writing the manuscript. All authors contributed to manuscript review and editing.

## Supporting information

Proteomics

## ACKNOWLEDGEMENTS

We thank all members of the Morrisey lab and the CHOP Division of Neonatology for their encouragement and support. We also thank the CHOP Flow Cytometry Core and the Penn CDB Microscopy Core for technical assistance. This research was supported by the National Institutes of Health (HL152194, HL132999, HL134745, HL148857 to E.E.M and 5T32HD043021 to D.A.C) and the

American Academy of Pediatrics Marshall Klaus Award (D.A.C.).

