## Supplemental for "TGFβ controls alveolar type 1 epithelial cell plasticity and alveolar matrisome gene transcription"

### SUPPLEMENTAL FIGURES

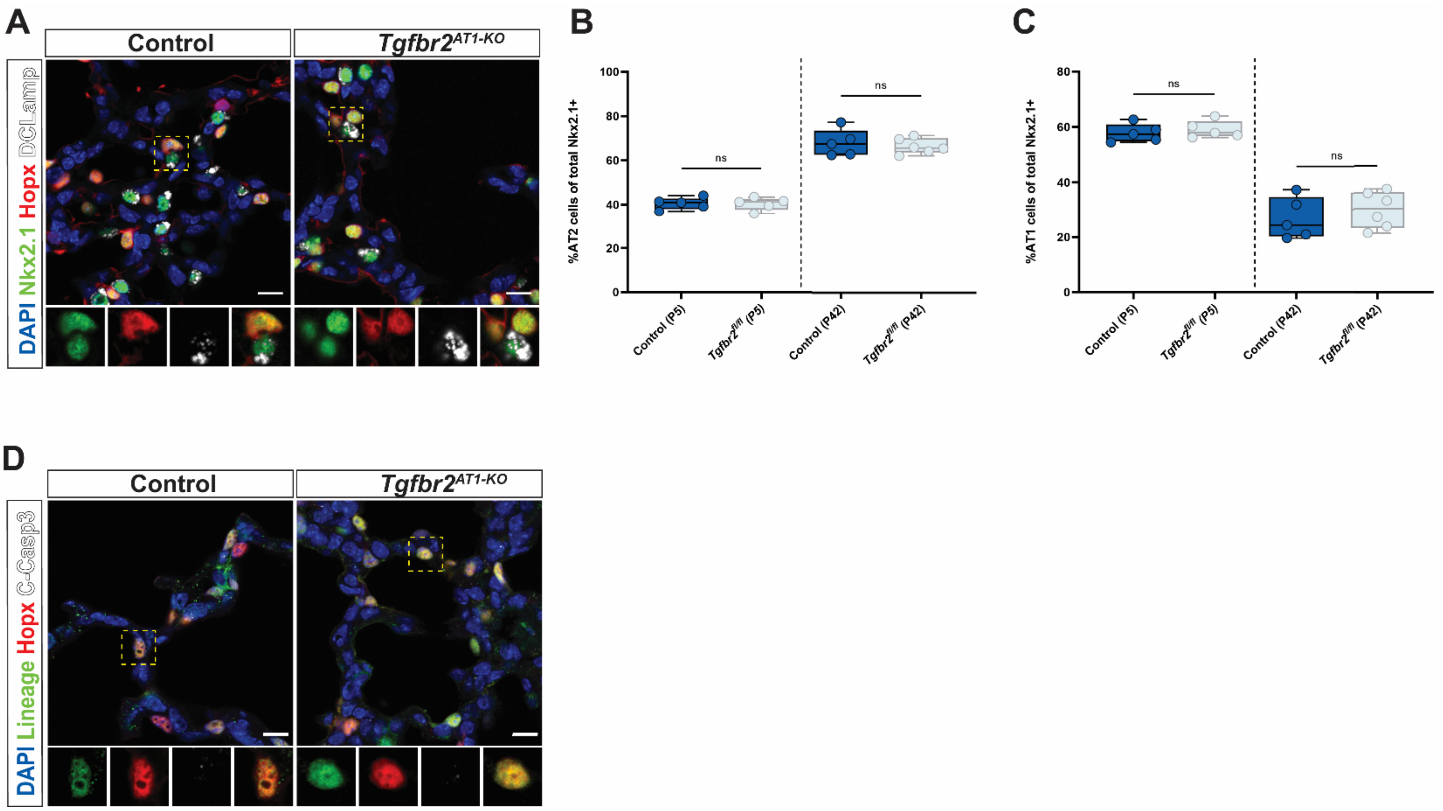


**Figure S1. AT1 cell reprogramming in Tgfbr2$^{AT1\text{-}KO}$ mice does not affect overall alveolar epithelial cell composition or apoptosis.**

A) IHC for Nkx2.1, HOPX, and DCLAMP demonstrate no change in total AT2 or AT1 cell numbers after postnatal loss of *Tgfbr2*. The yellow dashed box denotes the magnified region shown below the image and separated by fluorescence channel.

B) Quantification of AT2 cell numbers from total Nkx2.1+ cells in (A) denoting percent of cells that were Nkx2.1+ and DCLAMP+ at P5 (left) and P42 (right) by unpaired two-tailed t-tests with Welch's correction (n=5-6 per group).

C) Quantification of AT1 cell numbers from total Nkx2.1+ cells in (A) denoting percent of cells that were Nkx2.1+ and Hopx+ at P5 (left) and P42 (right) by unpaired two-tailed t-tests with Welch's correction (n=5-6 per group).

D) IHC for EYFP, HOPX, and cleaved caspase-3 do not indicate the presence of apoptotic cells after postnatal loss of *Tgfbr2*.

Each dot represents a single mouse. Scale bars denote 10 μm. ns = not significant. P values are denoted above the plots.

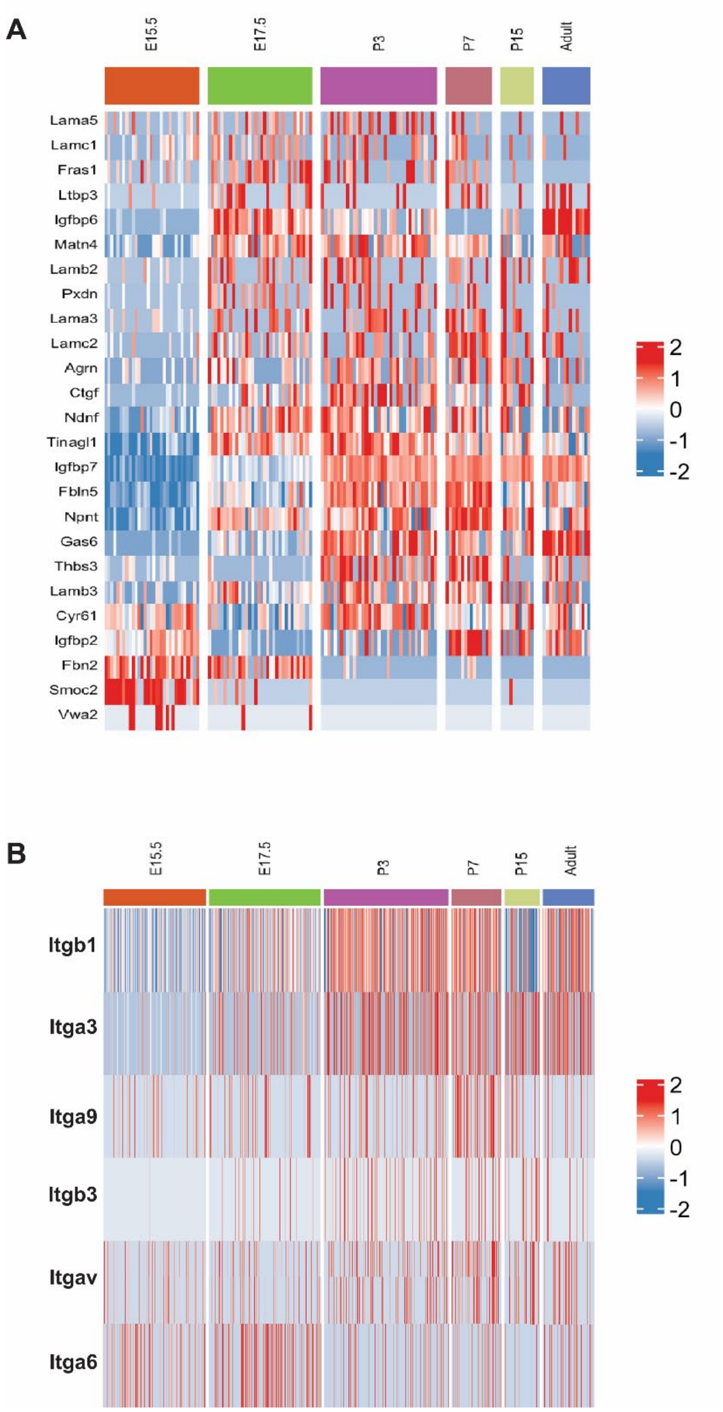


**Figure S2. AT1 cells exhibit a dynamic expression pattern for multiple matrisome and integrin genes starting in late lung development.**

A) Evaluation of previously generated scRNA-seq data across late lung development indicates that AT1-enriched glycoprotein and (B) integrin genes become highly expressed in the postnatal period indicating a role for AT1 cells in the active production and remodeling of the pulmonary matrisome across the lifespan.

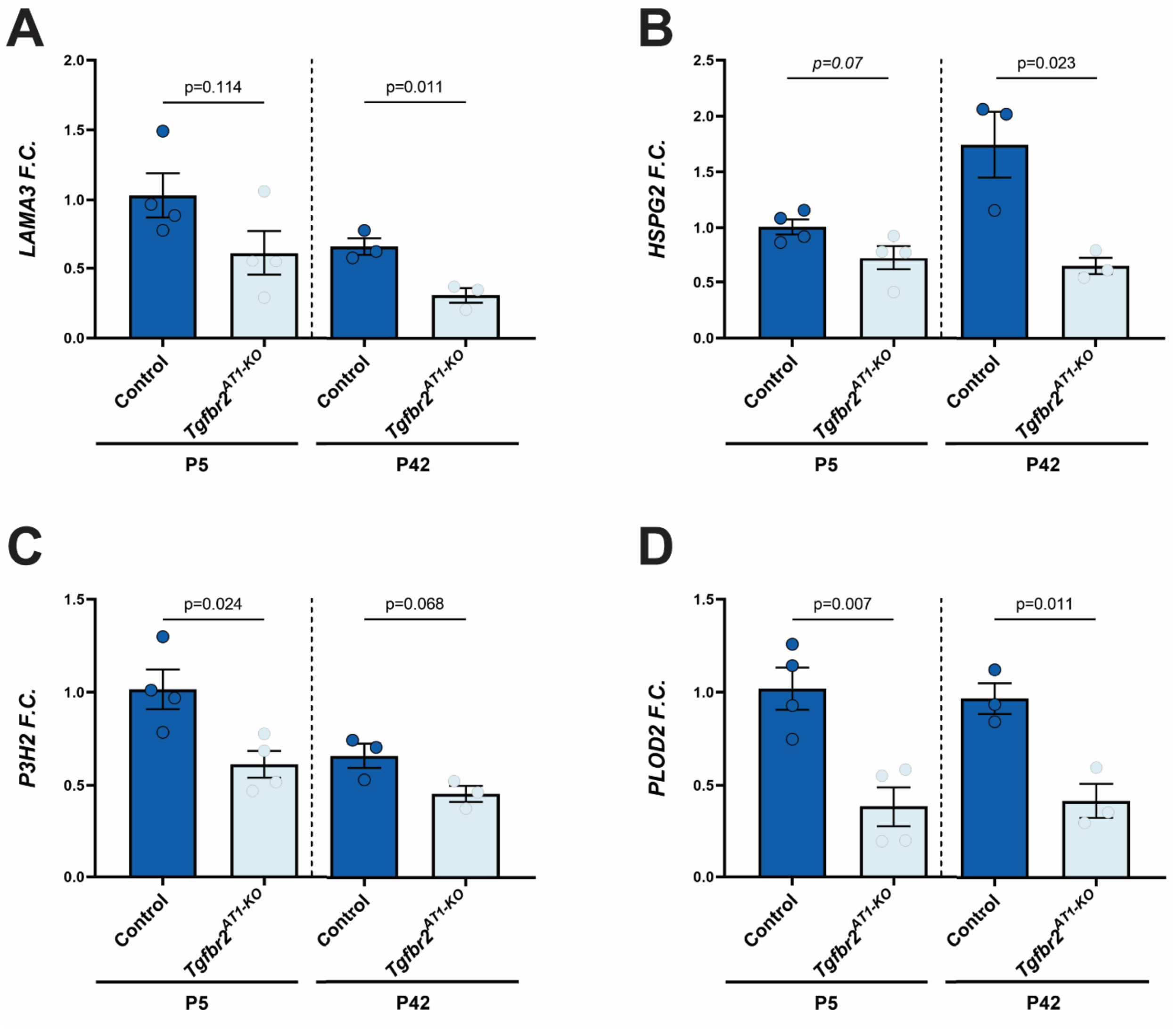


**Figure S3. Tgfbr2$^{AT1\text{-}KO}$ cells exhibit attenuated glycoprotein, proteoglycan, and matrisome regulatory enzyme expression**

A) *Tgfbr2*$^{AT1\text{-}KO}$ AT1 cells exhibit decreased RNA transcript expression for several AT1 cell-enriched core matrisome constituents including the glycoprotein (A) *Lama3,* the proteoglycan (B) *Hspg2*, and regulatory enzymes (C) *P3h2* and (D) *Plod2* at P5 that persists to P42 (n=3-4, unpaired two-tailed t-test with Welch's correction).

Each dot represents a single mouse. P values are denoted above the plots.
